# *Bipalium admarginatum* de Beauchamp, 1933 (Platyhelminthes, Tricladida, Geoplanidae) in Malaysia, with molecular characterisation including the mitogenome

**DOI:** 10.1101/2023.02.16.528827

**Authors:** Oi Yoon Michelle Soo, Romain Gastineau, George Verdon, Leigh Winsor, Jean-Lou Justine

## Abstract

We present here the first observation of *Bipalium admarginatum* de Beauchamp, 1933 since its original description 90 years ago. Three specimens were found on Perhentian Kecil Island, off Terengganu State, Malaysia and photographed in the field, and two were collected. This report thus includes the first colour photographs published for this species, from a locality close to the type-locality, Tioman Island (which is ca. 200 km south of the locality in this study, on the east coast of Peninsula Malaysia). We describe the external morphology and colour pattern of the species, which correspond well to the original description, itself based only on two preserved specimens. We performed an in-depth molecular characterisation of the species, including its complete mitochondrial genome, the 18S sequence and elongation 1 alpha sequence. In addition, EF1a sequences were also retrieved for 5 additional geoplanid species. No *tRNA-Thr* could be detected in the mitogenome of *B. admarginatum*, a lack already reported in several species of geoplanids, but we found a 13 bp sequence that contains the anticodon loop and seems to be conserved among geoplanids and might thus possibly represent a non-canonical undetected tRNA. We discuss the difficulties encountered in trying to reconstruct the cluster of nuclear ribosomal genes, a problem already mentioned for other Triclads. Three phylogenies, based respectively on all mitochondrial proteins, 18S, and EF1a, were computed; the position of *B. admarginatum* within the Bipaliinae was confirmed in each tree, as sister-group to various bipaliine species according to the sequences available for each tree.

## Introduction

Land flatworms (family Geoplanidae) include about 300 species, mainly found in tropical areas, and are famous for their ability to conquer new habitats worldwide and to become invasive alien species (Carbayo et al., 2016; Justine et al., 2021; Justine et al., 2022b; Justine et al., 2015; Justine et al., 2020b; Sluys, 2016; Winsor, 1983). The Bipaliinae or hammerhead flatworms are among the giants in this family, with species reaching almost one metre in length (Kawakatsu et al., 1982); their area of origin is South-East Asia and Madagascar. A recent study modelling potential distribution (Fourcade et al., 2022) found that the five more common bipaliines, already found on various continents, could still invade more regions of the world. According to scenarios of future climate change, two species (*Bipalium kewense* Moseley, 1878 and *B. vagum* Jones & Sterrer, 2005) that already have the largest observed global range are predicted to further increase their potential distribution (Fourcade et al., 2022).

This study is not about an invasive species found far away from its area of origin, but instead it is about a rarely recorded species, *Bipalium admarginatum* de Beauchamp, 1933, which we collected in the area where it was first described, islands off the East coast of Peninsular Malaysia. We subjected our specimens to molecular techniques that had been, until now, mostly used on invasive species. The complete mitogenome was sequenced and compared with existing data (Gastineau et al., 2019; Justine et al., 2022a). The phylogenetic position of *B. admarginatum* was studied with 3 different datasets. Special attention was paid to the search for a missing mitochondrial tRNA. Finally, an attempt to reconstruct the two variants of the cluster of nuclear ribosomal RNA genes was undertaken.

This paper provides information about a rarely mentioned species and molecular data which we hope will be useful for comparison with other geoplanids.

## Material and methods

### Origin of the specimens

Observations, photographs, and collections were performed by one of us (GV). Three specimens were found on the soil on the edge of primary jungle on the east coast of Perhentian Kecil Island, off Terengganu State in Peninsular Malaysia. The first one, found on 26 August 2019, was photographed in natural light but not collected. Two specimens were found (5°54’ 22.2”N 102°43’ 55.7”E) on 28 September 2019; they were photographed live (both in natural light and under artificial light) and then fixed in Gin (ca. 40% ethanol), in the absence of a more scientific fixative. After several days, the specimens were shipped to Kuala Lumpur for examination, and the Gin was replaced by 100% ethanol. Specimens were then shipped to Paris and deposited in the collection of the Muséum National d’ Histoire Naturelle under registration numbers MNHN JL354 and MNHN JL355. Photographs of preserved specimens were taken, and the posterior part of the body of specimen MNHN JL354, about 1 centimetre in length, was taken for molecular studies.

### Morphometric data

Morphometric data of living and preserved specimens were calculated from scaled photographs. Colour names and codes were taken from the RAL colour chart at https://www.ralcolorchart.com/.

### Sequencing and assembly

A piece of specimen MNHN JL354 of *B. admarginatum* was sent to the Beijing Genomics Institute (BGI) in Shenzhen (China). DNA was extracted by BGI in accordance with their internal protocol. A total of ca. 80M 150 bp paired-end reads were obtained and assembled using SPAdes 3.15.5 (Bankevich et al., 2012) with a k-mer parameter of 125. The sequences of interest were found among the contig file resulting from assembly by using blastn command-line (Boratyn et al., 2012) with databases made of the full mitogenome, partial 18S and partial elongation factor 1-alpha genes of *Bipalium adventitium* Hyman, 1943 (GenBank accession numbers MZ561467, MZ520993 and KJ599681, respectively). Annotation of the mitogenome was carried out using MITOS (Bernt et al., 2013). The position of tRNA was checked using Arwen v1.2 (Laslett & Canbäck, 2008). The OGDRAW web portal was used to draw the map of the mitogenome (Lohse et al., 2013). The LOGO figure of the tRNA-Thr alignment was obtained using WebLogo3 online (Crooks et al., 2004).

### Phylogeny

Three different maximum likelihood phylogenies were generated. The first two were obtained by appending the new data on *B. marginatum* to datasets already used in previous papers (Gastineau et al., 2022; Justine et al., 2022a), consisting of concatenated alignments of the mitochondrial proteins and alignment of the partial 18S gene. The third one was obtained from alignment of the partial elongation factor 1-alpha (EF1-alpha) gene obtained in the course of this study with references coming from previous works (Almeida et al., 2021; Álvarez-Presas & Riutort, 2014; Carbayo et al., 2013) for a total of 279 sequences. The percentage identity between EF1a sequences was calculated after alignment by Clustal Omega (Sievers et al., 2011). All other alignments were done using MAFFT 7 (Katoh & Standley, 2013) with the -auto option. For the multigene phylogeny, amino-acid sequences were aligned separately, trimmed and then concatenated using Phyutility 2.7.1 (Smith & Dunn, 2008). All trimming was performed with trimAl (Capella-Gutiérrez et al., 2009) and the - automated1 option. Evolutionary models were selected based on results returned by ModelTest-NG v0.1.7 (Darriba et al., 2019), with the concatenated sequence of mitochondrial proteins being considered a single amino-acid sequence. Maximum likelihood phylogenies were obtained using IQ-TREE 2.2.0 (Minh et al., 2020) after 1 000 bootstrap replications.

## Results

### Description of the specimens (Figures 1-3)

An elongate living specimen (MNHN JL355) measured approximately 103 mm long, and 2.9 mm maximum wide over the pharyngeal region (**Figure 2B**). The fan-shaped headplate measured 2.1 mm in maximum width, less than the maximum body width, and had a length to width ratio of 1:1.8, with very slightly recurved small lappets. The transverse body shape is broadly convex. Eyes are visible as fine black spots, mainly around the lateral quarters of the headplate, and continuing posteriorly just above the marginal black stripes.

**Figure 1.**
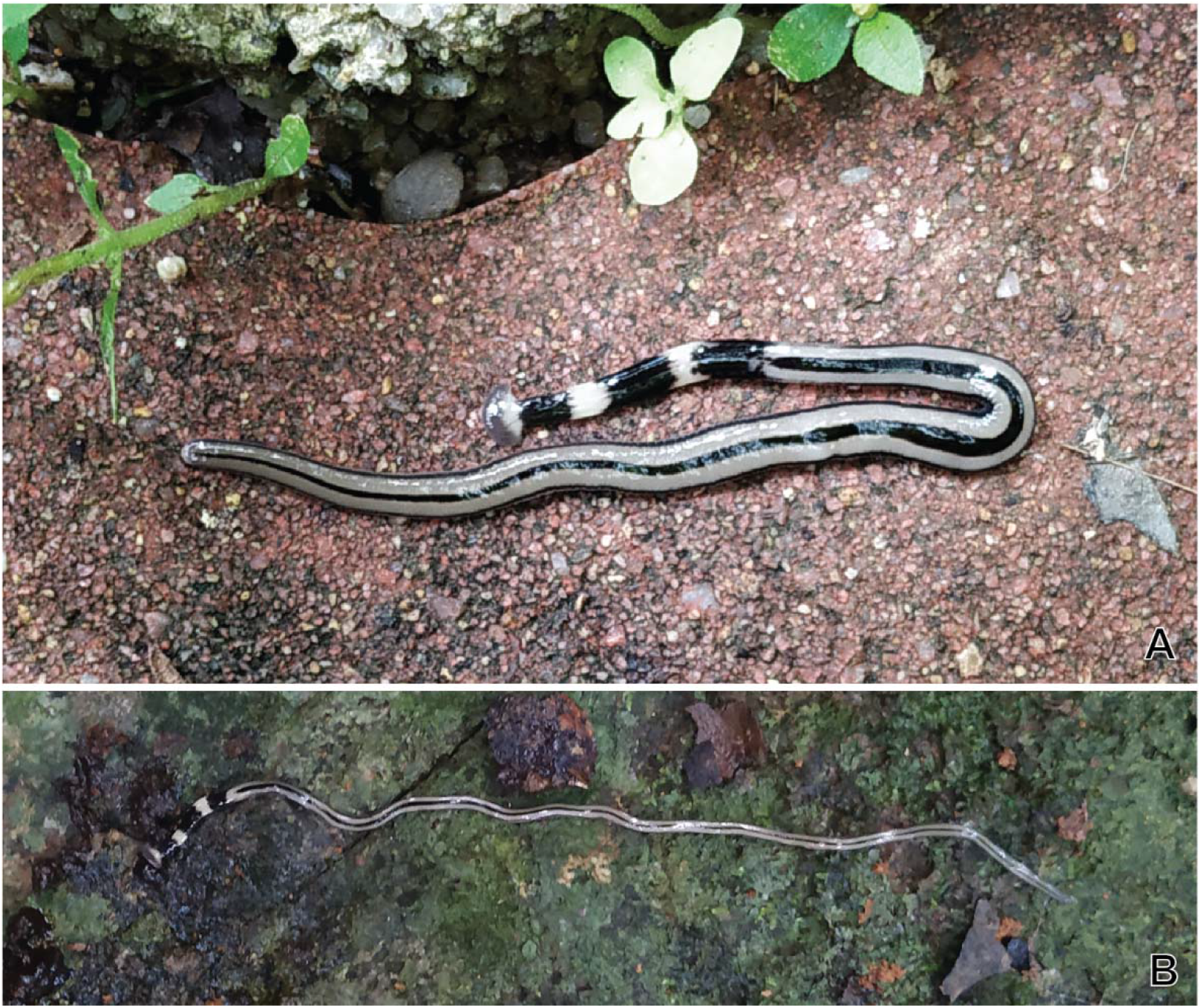
*Bipalium admarginatum*, live specimens photographed in the field under natural light. A, specimen not collected, photographed 16-08-2019. B, specimen MNHN JL354 (a part of this specimen was used for the molecular analysis). Unscaled. Photos by George Verdon.

**Figure 2.**
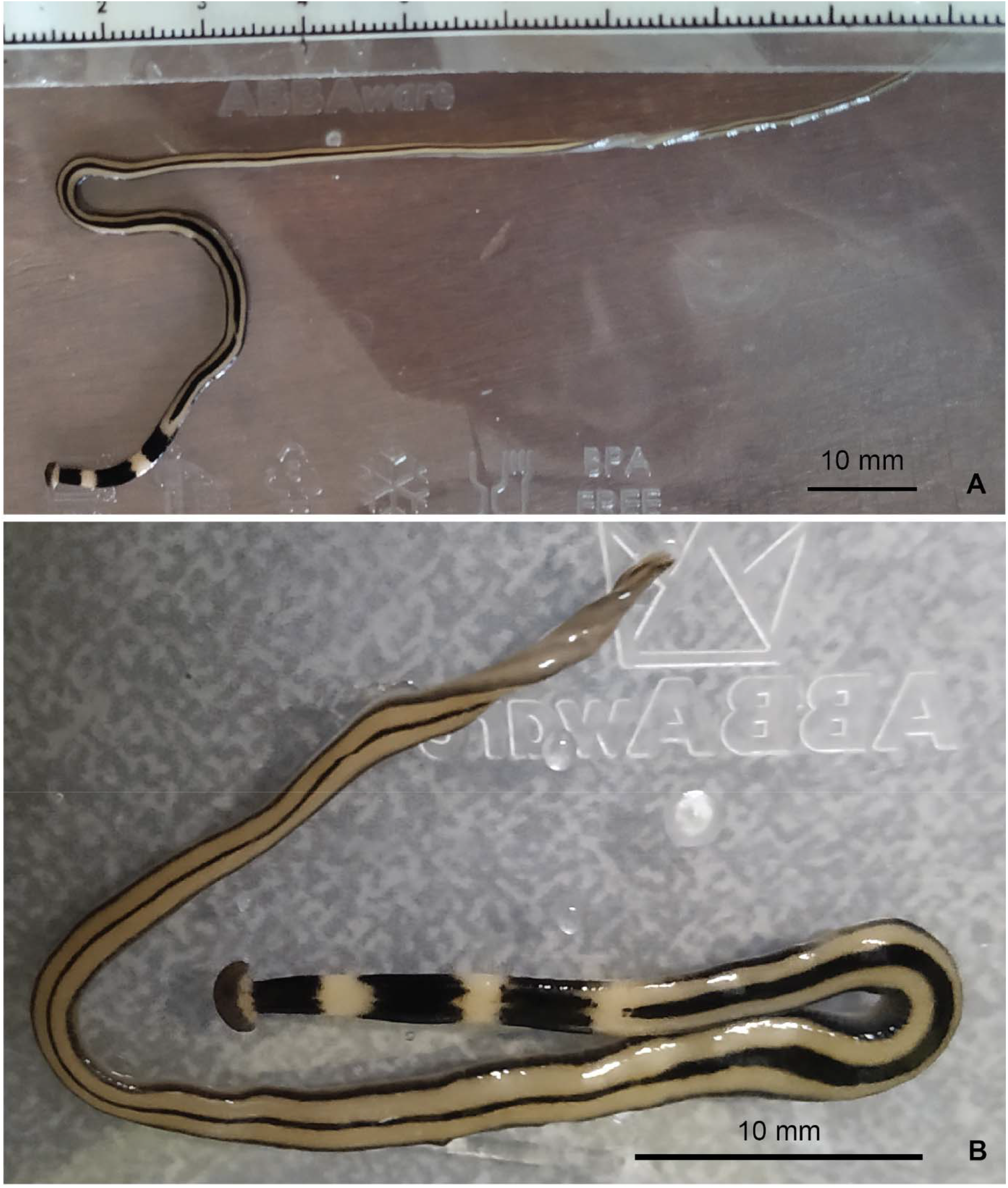
*Bipalium admarginatum*, live specimens photographed under artificial light. A, specimen MNHN JL354; B, Specimen MNHN JL355. Photos by George Verdon.

The preserved specimen (MNHN JL355) measured 77.7 mm long, and 4.1 mm maximum wide over the pharynx, with the mouth 25.0 mm from the anterior end, and gonopore 5.9 mm behind the mouth (**Figure 3B**). The narrow creeping sole is slightly raised above the convex surface.

**Figure 3.**
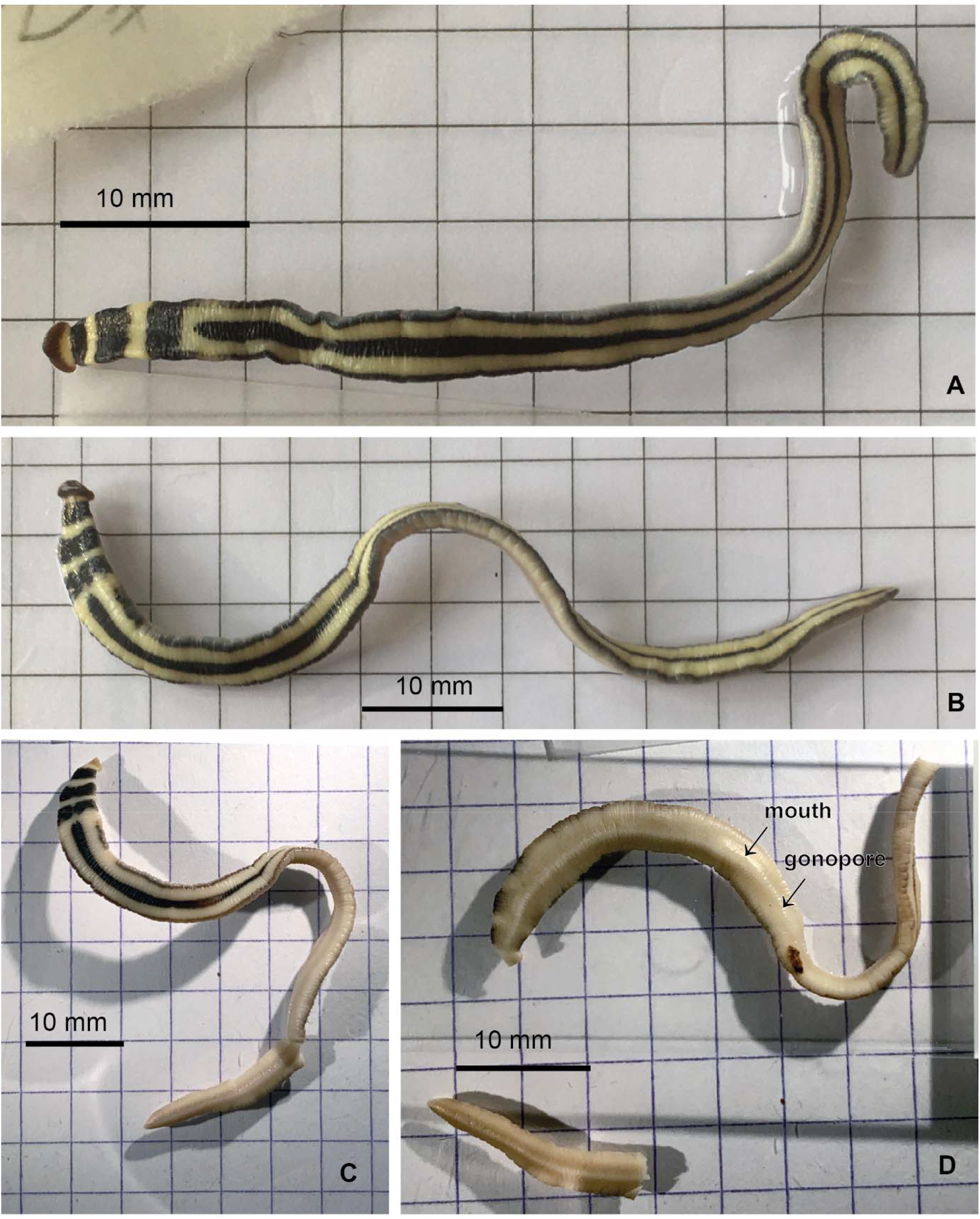
*Bipalium admarginatum*, preserved specimens. A, specimen MNHN JL354, undamaged; B-D, specimen MNHN JL355; B, undamaged; C, partially showing ventral surface; D, damaged, showing ventral surface and position of mouth and gonopore. A, B, photographs taken in 2019 vs. C, D, photographs taken in 2023, note that yellow colour has vanished. Photos by Jean-Lou Justine.

Since colours appear slightly different for specimens photographed in natural light and under artificial light, we show photographs taken under both conditions (**Figures 1-2**). The colour and pattern of the living specimen (**Figure 1**) are as follows: the anterior two-thirds of the headplate, reaching in a curve from lappet to lappet, is Black-grey (RAL 7021) in colour, with the posteriad headplate lighter Telegrey 4 (RAL 7047). Behind the headplate are three transverse bands, Jet Black (RAL 9005) in colour. The margins of these bands are irregular, and the length of each band is slightly greater than the one before it. The black bands are separated by two transverse bands coloured Traffic White (RAL 9016). The dorsal ground colour of the 85% of the remaining body is slightly darker than Telegrey 4. Three dorsal longitudinal stripes that terminate just short of the rounded posterior tip are present. They comprise a broad, gently tapering mid-dorsal Jet Black stripe extending from just behind the hindmost black band, and widest over the pharyngeal region, and paired narrow Jet Black marginal stripes of even thickness that extend posteriorly from the last black transverse band. In the living specimen, a fine faint grey, discontinuous longitudinal stripe, slightly darker than the ground colour, is present between the median and marginal stripes, but not visible in the preserved specimen. The ventral ground colour is Telegrey 4, with whitish creeping sole delineated on either side by a fine Black-grey longitudinal stripe. In the anterior ventral surface of the preserved specimen (JL355), the two white bands, and the first two transverse black bands, though less strongly pigmented, wrap around the body as far as the ventral zone.

The form of the dorsal transverse bands and longitudinal stripes appeared constant in the other specimens examined (JL354), as well as a specimen from Tioman Island Mersing, Jahor, Malaysia (iNaturalist 21 February 2021) https://www.inaturalist.org/observations/69891718 *accessed January 2023*). The ventrad extension of the transverse bands was observed in both preserved specimens JL354 and JL355. Morphometric data for the preserved specimens are provided in **Table 1**. Neither of these specimens were examined histologically.

**Table 1.** The three specimens found and their measurements. Percentages are of total length

| Specimen | Living |  |  | Preserved |  |  |  |
| --- | --- | --- | --- | --- | --- | --- | --- |
|  | Length | Length | Width | Distance of the mouth from the anterior end |  | Distance of the gonopore from the mouth |  |
| Unit | mm | mm | mm | mm | % | mm | % |
| Not collected | ca. 45 | - | - | - | - | - | - |
| MNHN JL354 | ca. 131 | 58.0 | 3.6 | 15.9 | 27.4 | - | - |
| MNHN JL355 | ca. 103 | 77.7 | 4.1 | 25.0 | 32.2 | 5.9 | 6.9 |
| de Beauchamp, 1933 | - | 85 | 3 | 30 | 35.3 | 6 | 7.1 |

### Mitogenome

Although the specimen was originally processed with a somewhat unusual protocol, including fixation in a non-scientific ethanol solution (Gin), sequencing allowed to recover the mitogenome with a high coverage.

A 19 115 bp long contig with a coverage of 178.61 X and redundant endings was retrieved from the assembly. After trimming and circularisation, the mitogenome (GenBank OQ308795) is 18 990 bp long **(Figure 4)**. It codes for 12 conserved protein-coding genes, 21 tRNA and 2 rRNA. It is for now the largest known mitogenome for a Bipaliinae (Table 2). This extra length results from intergenic sequences, especially a large region located between *rrnL* and *cob*. This region is 2 288 bp long and contains only 2 tRNA. A comparison of the size of this region among Bipaliinae is presented in **Table 2**. In a more general way, it should be noted that the mitogenome is not compact, with intergenic lengths parsed among it.

**Figure 4.**
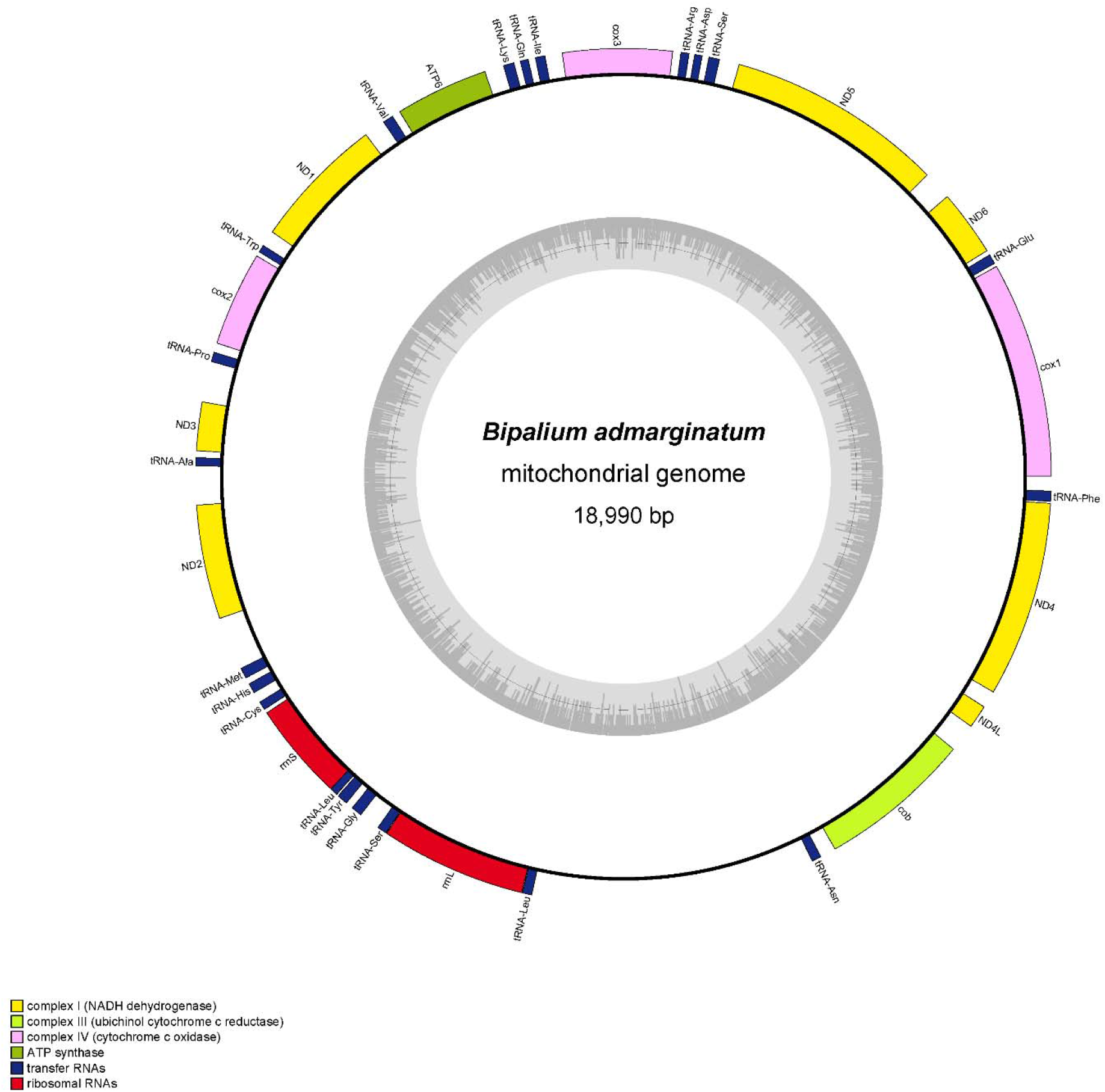
*Bipalium admarginatum*, map of mitochondrial genome. The mitogenome codes for 12 conserved protein-coding genes, 21 tRNA and 2 rRNA.

**Table 2.** Mitogenomes of Bipaliinae and various characteristics

| Species | GenBank | Size | Length of the *rrnL-cob* intergenic region and presence of tRNA |
| --- | --- | --- | --- |
| *Bipalium admarginatum* | OQ308795 | 18 990 bp | 2288 bp (*tRNA-Leu*, *tRNA-Asn*) |
| *Bipalium kewense* | MK455837 | 15 666 bp | 934 bp (*tRNA-Leu*, *tRNA-Thr*, *tRNA-Asn*) |
| *Bipalium vagum* | MZ561468 | 17 149 bp | 1552 bp (*tRNA-Leu*, *tRNA-Thr*, *tRNA-Asn*) |
| *Bipalium adventitium* | MZ561467 | 15 494 bp | 741 bp (no tRNA detected) |
| *Diversibipalium multilineatum* | MZ561469 | 15 660 bp | Impossible to circularize (*tRNA-Leu*, *tRNA-Thr*, *tRNA-Asn*) |
| *Diversibipalium mayottensis* | MZ561470 | 15 989 bp | 1479 bp (*tRNA-Leu*, *tRNA-Thr*, *tRNA-Asn*) |
| *Humbertium covidum* JL351 | MZ561472 | 15 540 bp | 691 bp (*tRNA-Leu*, *tRNA-Asn*) |
| *Humbertium covidum* JL090 | MZ561471 | 15 524 bp | 672 (*tRNA-Leu*, *tRNA-Asn*) |

### The case of the missing tRNA-Thr

No tRNA-Thr could be detected in the mitogenome of *B. admarginatum*. This lack has already been noticed in the Bipaliinae *Humbertium covidum* Justine, Gastineau, Gros, Gey, Ruzzier, Charles & Winsor, 2022 (Justine et al., 2022a), but also among three species of Rhynchodeminae, namely *Parakontikia ventrolineata* (Dendy, 1892) Winsor 1991 (Gastineau & Justine, 2020), *Platydemus manokwari* de Beauchamp, 1963 (Gastineau et al., 2020) and *Australopacifica atrata* (Steel, 1897) (Gastineau et al., 2022).

We tried to align the *tRNA-Thr* genes from mitogenomes of Geoplanidae where they were detected, including the Geoplaninae *Obama nungara* Carbayo, Álvarez-Presas, Jones & Riutort, 2016 (Solà et al., 2015) and *Amaga expatria* Jones & Sterrer, 2005 (Justine et al., 2020a), with the complete mitogenomic sequences of *B. admarginatum* and *H. covidum* JL090 and JL351. The *tRNA-Thr* genes aligned in an area that corresponds to their assessed position in other species, between *cob* and *rrnl*. The alignment showed the presence of a nearly perfectly conserved 13 bp sequence that contains the anticodon loop as it can be seen on the LOGOS alignment provided as **Figure 5**. Only *Diversibipalium mayottensis* Justine, Gastineau, Gros, Gey, Ruzzier, Charles & Winsor, 2022 and *Bipalium vagum* exhibited each a single polymorphism in this sequence. The 5’ and 3’ ends of the 72 bp sequence derived from *B. admarginatum* were completely conserved for 4 bp in 5’ and 6 bp in 3’ with *Bipalium kewense* Moseley, 1878. Attempting to manually model a functional tRNA for *B. admarginatum* based on this sequence remained non-conclusive (figure not shown), leading to a tRNA that would have the following characteristics: an acceptor stem with a single mismatch out of 7 pairs, but one mismatch plus one unpaired base in the anticodon arm; complete lack of a D-arm.; the putative TΨC-arm would have had at maximum 3 well matched pairs of nucleotides and 21 unpaired bases.

**Figure 5.**
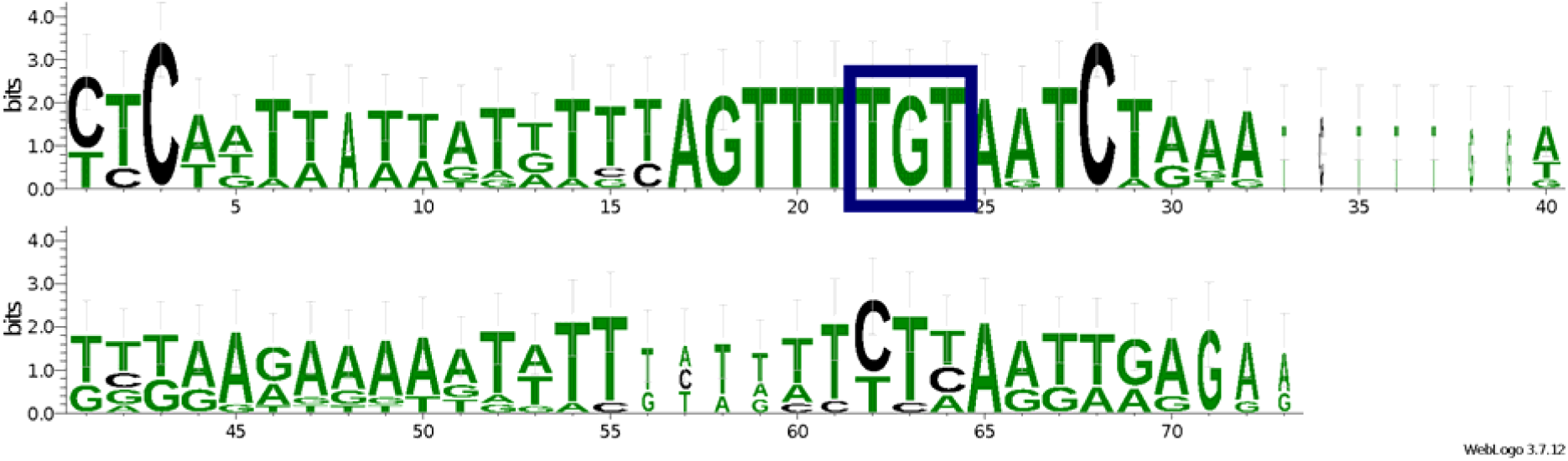
LOGOS alignment of the *tRNA-Thr* genes from mitogenomes of geoplanids.

### Trying to reconstruct the cluster of nuclear ribosomal RNA genes

The cluster of rRNA did not come as a single sequence of high coverage. Instead, several contigs returned a positive blastn result. These contigs varied in both size and coverage. **Table 3** summarizes the names of the contigs that were used in our attempt to reconstruct the rRNA, with their sizes and coverages.

**Table 3.**
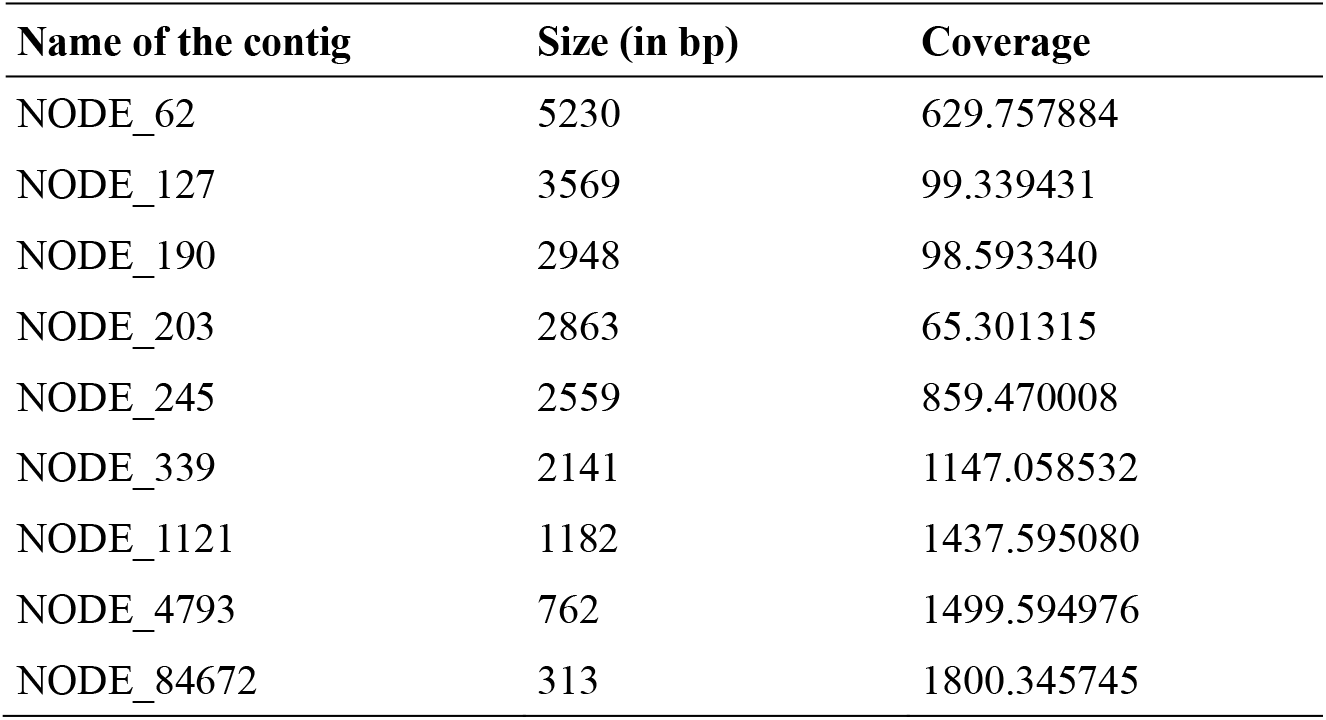
Contigs used to reconstruct the nuclear ribosomal RNA genes

Some of these contigs were overlapping with others by a number of 125 bp, corresponding to the k-mer used for assembly. Two of these highly covered contigs, NODE_4793 and NODE_84672, were found to merge with both high and low coverages contigs. This situation was understood as the presence of two types of rRNA, first evidenced among Dugesiidae (Carranza et al., 1999; Carranza et al., 1996) and later among other Tricladida (Breugelmans et al., 2012). We attempted to reconstruct the complete clusters based on the coverage of the largest contigs, which we will therefore refer to as ‘ high’ and ‘ low’. **Table 4** provides the names and order of the contigs used in the attempt to reconstruct both clusters.

**Table 4.** Contigs used in the attempt to reconstruct both clusters

| Type | Contigs (5' to 3', starting with 18S) |
| --- | --- |
| High | NODE_62-NODE_4793-NODE_1121-NODE_339-NODE_84672-NODE_245 |
| Low | NODE_203-NODE_4793-NODE_190-NODE_84672-NODE_127 |

For the SSU phylogeny computed in this paper, only contig NODE_4793 was used, as it is common to both putative types and also corresponds to the portion of the 18S gene previously used in Justine *et al*. (2022). Only this fragment has been deposited in GenBank (accession number: OQ308840). The sequences of the two putative clusters can be downloaded from Zenodo, as indicated in the Data availability statement.

### EF1-alpha gene

It was possible to retrieve the complete EF1-alpha gene of *B. admarginatum* (OQ326523). The intronless gene is 1 392 bp long. This complete coding sequence was later used to datamine the previous sequencing results from Gastineau *et al*. (2019) and Justine *et al*. (2022) (Gastineau et al., 2022; Justine et al., 2022a). It was possible to retrieve the same complete coding sequence for *Bipalium kewense* JL184A (OQ326519), *Humbertium covidum* JL090 (OQ326520), *Diversibipalium multilineatum* JL177 (OQ326521) and *Bipalium vagum* JL307 (OQ326522). All genes were of identical lengths. **Table 5** lists the species, the GenBank accession number of their EF1-alpha gene obtained in the course of this study. and the percentage of identity between them as calculated after alignment by Clustal Omega.

**Table 5.** Species of Bipaliinae and the percentage identity of EF1a gene between them

| Name | *B. vagum* | *H. covidum* | *B. admarginatum* | *B. kewense* | *D. multilineatum* | GenBank |
| --- | --- | --- | --- | --- | --- | --- |
| *B. vagum* | 100.00 | 90.23 | 91.02 | 91.09 | 91.24 | OQ326522 |
| *H. covidum* |  | 100.00 | 91.88 | 91.95 | 92.60 | OQ326520 |
| *B. admarginatum* |  |  | 100.00 | 94.32 | 94.47 | OQ326523 |
| *B. kewense* |  |  |  | 100.00 | 97.92 | OQ326519 |
| *D. multilineatum* |  |  |  |  | 100.00 | OQ326521 |

### Molecular phylogenies

The best models of evolution suggested by ModelTest-NG were mtZOA+I+G4 for the mitochondrial proteins, TVM+I+G4 for 18S and GTR+I+G4 for EF1-alpha. The two first phylogenies were congruent with previously published works (Justine et al., 2022). The amino acid inferred phylogeny (**Figure 6**) placed *B. admarginatum* as sister species to a clade containing *B. kewense* and *Diversibipalium multilineatum* Makino & Shirasawa, 1983, with strong support. In the main clade of the Geoplanidae, the lowest support was found at the node separating Bipaliinae and Rhynchodeminae, with only 66% support. The SSU-inferred phylogeny (**Figure 7**) associated *B. admarginatum* with *Novibipalium venosum* (Kaburaki, 1922), but with weak support at the node. For the EF1-alpha inferred tree, only the subtree containing the Bipaliinae is shown (**Figure 8**). The tree associated *B. admarginatum* with *Bipalium adventitium*, although with low support at the node. The tree displayed *B. vagum* at some distance from the rest of the Bipaliinae in a similar way to the mitochondrial protein inferred phylogeny.

**Figure 6.**
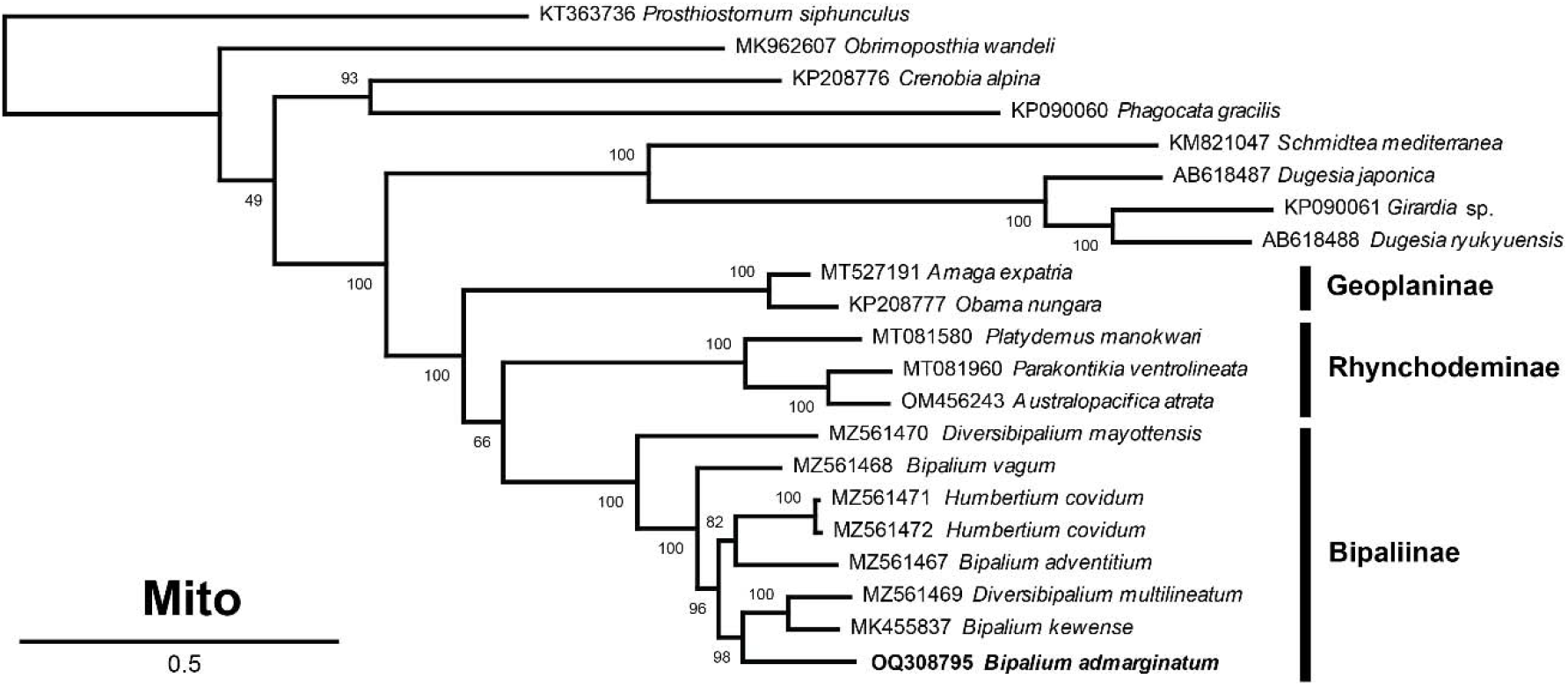
Maximum likelihood phylogenetic tree (mtZOA+I+G4 model) obtained from concatenated amino acid sequences of the mitochondrial proteins of *Bipalium admarginatum* and other flatworms. The tree with the best likelihood is shown and ML bootstrap support values are indicated. Subfamilies of Geoplanidae are indicated on the right. Based on the matrix and method used in Gastineau et al. (2022) with addition of the new sequence of *B. admarginatum*.

**Figure 7.**
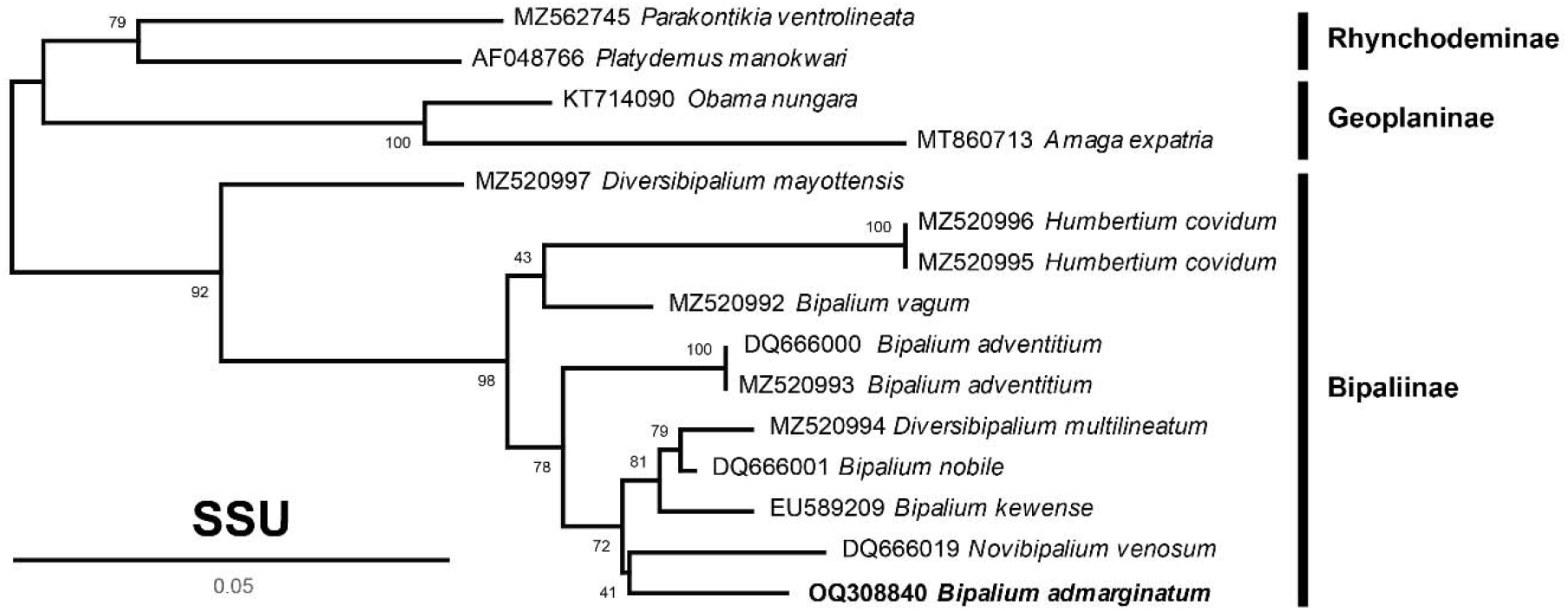
SSU phylogenetic tree of bipaliine geoplanids. Maximum likelihood phylogenetic tree based on 15 partial SSU genes, using the TVM+I+G4 model of evolution. The tree with the best likelihood is shown, and ML bootstrap support values are indicated. The subfamilies within the Geoplanidae (Rhynchodeminae, Geoplaninae and Bipaliinae) are indicated. Based on the matrix and method in Justine et al. (2022) with the addition of the new sequence of *Bipalium admarginatum*.

**Figure 8.**
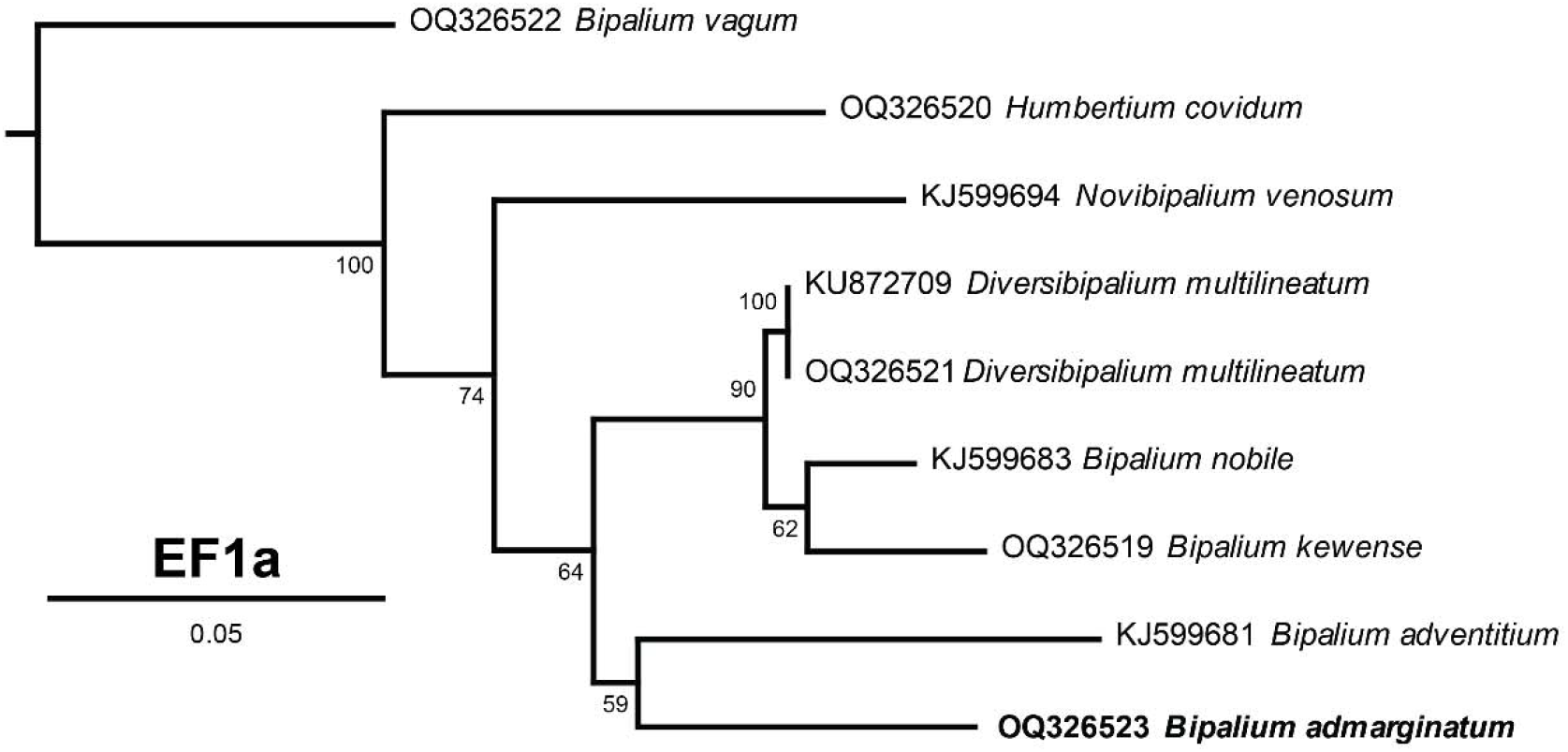
Phylogenetic tree of geoplanids based on nine sequences of elongation factor-1 alpha, including five new; only the part of the tree containing the Bipaliinae is shown. Maximum likelihood phylogenetic tree based on 9 sequences, using the GTR+I+G4 model of evolution; all OQ sequences are our own and new.

## Discussion

### Identification and rediscovery of *Bipalium admarginatum*

*Bipalium admarginatum* was described by de Beauchamp (1933) from two specimens collected in Sedagong, altitude 1 000 feet, on Tioman Island, in 1927. The illustration associated with the description includes a schematic diagram of the eyes on the head and a longitudinal section of the histology of the copulatory apparatus, and a half-tone drawing of the dorsal side of what is approximately the anterior half of a specimen.

The locality where we found our specimens (Perhentian Kecil Island) is ca. 200 km north of the type-locality (Tioman Island) and both localities are on the east coast of Peninsular Malaysia. Interestingly, each record was on an island. There is a single observation of this species in iNaturalist, in Tioman Island, the type-locality, on 12 May 2018 (iNaturalist observation #69891718). The present paper is the first scientific record of the species since its original description in 1933, 90 years ago.

We present here the first coloured photograph of a living specimen of *Bipalium admarginatum*. The ground colours of our specimens differ significantly from the light brown of the preserved specimens from Tioman Island upon which the original description (De Beauchamp, 1933) was based. The brown colour of these specimens was probably artefactual alcohol browning. Otherwise, the band, stripe, and eye pattern of the specimens examined in this study agree closely with the original description. Our preserved specimens are not as long as the two examined by de Beauchamp (85 mm and 90 mm), but the relative positions of the mouth and gonopore of our specimen (JL355) align well with those in the original description (**Table 1**). The extension of the transverse bands to the ventral zone, observed in both preserved specimens of *B. admarginatum*, has also been observed in another transversely and longitudinally striped species, *Diversibipalium boehmigi* (Müller, 1902) recorded from Mount Matang, Sarawak, Malaysia (Müller, 1902).

### Molecular characteristics

The fact that we found a conserved pattern of the anticodon loop of the *tRNA-Thr* among *B. admarginatum* and *H. covidum* is intriguing at least, keeping in mind that all dedicated software failed until now to detect it among these mitogenomes. We are not ruling out the possibility that this tRNA indeed exists with a heavily modified structure as it can sometimes be found among Metazoa (Krahn et al., 2020). Armless mitochondrial tRNA have, for example, been found *in silico* among Nematoda (Jühling et al., 2012; Jühling et al., 2018) and their presence assessed by biochemical proofs (Wende et al., 2014). However, in the absence of similar proofs, we refrain from including this *tRNA-Thr* in our annotations, which does not preclude looking for the conserved anticodon loop among other mitogenomes of Geoplanidae among which this tRNA was not detected.

In a comparable way, we have chosen to restrict our use of the nuclear rRNA to a portion of the 18S gene that seems to be common to the two types, and only submitted this fragment to GenBank. Although our ‘ hand-made’ reconstruction of these two clusters is plausible, we are concerned that we have possibly reached the boundaries of what can be obtained by short-read sequencing technologies. Perhaps ‘ long-read’ technologies such as those that sequence native DNA will prove fruitful in the near future. We are nonetheless convinced of the importance of this topic for its evolutive significance, the molecular and biochemical implications it induces, but also because of the putative bias it can induce in phylogeny and molecular taxonomy.

## Conclusion

This study is probably the first of a complete mitogenome for a geoplanid which has *not* been recorded as an invasive species. We expect to continue such studies on various members of the geoplanids, which will provide additional data to understand the evolution of land flatworms.

This study is also pioneering with regard to the determination of EF1a sequences; it will be necessary to confirm on other species whether these sequences are interesting in terms of understanding the phylogeny of the Geoplanidae.

Regarding conservation issues, we note that this species has only been found on islands off the east coast of Peninsular Malaysia, which are probably more unspoiled habitats than the coast. It would be interesting to look for this species, and others, on the mainland.

We also note that excellent sequencing results were obtained for specimens originally preserved in the field with a very non-scientific product (Gin), pending the correct fixation of the specimens in the laboratory in 95% ethanol. This is encouraging for future research. In the absence of the proper DNA fixative, those who unexpectedly happen upon interesting species could temporarily preserve the specimens in a similar way to that described here, until the flatworms can reach a laboratory. We already have molecular results (unpublished) on a flatworm which was fixed in locally produced moonshine.

## Data availability

All the sequences obtained in the course of this study can be downloaded from Zenodo following this link: https://doi.org/10.5281/zenodo.7573203

## Acknowledgements

We thank Dr Tan Koh Siang from the Raffles Museum who kindly checked whether any colour illustration of *Bipalium admarginatum* was available and provided documentation. We also thank Liv Grant for assistance in the field; this included fending off monkeys during flatworm collection.

## Notes

https://doi.org/10.5281/zenodo.7573203

### Competing Interest Statement

The authors have declared no competing interest.

